# iHDSel software: The Price equation and the population stability index to detect genomic patterns compatible with selective sweeps. An example with SARS-CoV-2

**DOI:** 10.1101/2024.06.01.596790

**Authors:** Antonio Carvajal-Rodríguez

## Abstract

A large number of methods have been developed and continue to be developed for detecting the signatures of selective sweeps in genomes. Significant advances have been made, including the combination of different statistical strategies and the incorporation of artificial intelligence (machine learning) methods. Despite these advances, several common problems persist, such as the unknown null distribution of the statistics used, necessitating simulations and resampling to assign significance to the statistics. Additionally, it is not always clear how deviations from the specific assumptions of each method might affect the results.

In this work, allelic classes of haplotypes are used along with the informational interpretation of the Price equation to design a statistic with a known distribution that can detect genomic patterns caused by selective sweeps. The statistic consists of Jeffreys divergence, also known as the population stability index, applied to the distribution of allelic classes of haplotypes in two samples. Results with simulated data show optimal performance of the statistic in detecting divergent selection. Analysis of real SARS-CoV-2 genome data also shows that some of the sites playing key roles in the virus’s fitness and immune escape capability are detected by the method.

The new statistic, called *J*_*HAC*_, is incorporated into the iHDSel (informed HacDivSel) software available at https://acraaj.webs.uvigo.es/iHDSel.html.

## Introduction

Evolutionary biology studies the factors that affect genetic variability in populations and species. The main processes that influence the evolution of this variability include mutation and recombination, genetic drift, migration, and natural selection. Natural selection, in addition to affecting the allele carrying a beneficial mutation, impacts the neutral alleles of loci linked to the selective one, producing what is known as genetic hitchhiking (Smith and Haigh 1974; Kaplan et al. 1989), which leads to a selective sweep (Berry et al. 1991; Stephan 2019), meaning a loss of diversity around the selected site. These sweeps can be complete or incomplete, strong or soft, and they can even overlap (Johri, Stephan, et al. 2022). Regarding the detection of the footprint left by selective sweeps in genomes, from the earliest methods that explored haplotype patterns, whether by studying homozygosity (Sabeti et al. 2007), its diversity (Kimura et al. 2007), or interpopulation differentiation (Chen et al. 2010), among others, a great number of methods have been developed and continue to be developed. Significant advancements have been made, including the use of summary statistics, the combination of different statistical strategies, and the incorporation of artificial intelligence-based methods (Horscroft et al. 2019; Stephan 2019; Abondio et al. 2022; Arnab et al. 2023; Panigrahi et al. 2023; Whitehouse and Schrider 2023).

Most methods for detecting selective sweeps require the existence of haplotypic data, although see (Kern and Schrider 2018) where summary statistics calculated on unphased genotypes are used, which in a supervised machine learning context allow the classification of genomic windows subject to selection. However, supervised methods are computationally expensive and are highly dependent on training data, and their performance with data from other species, genome types and in general, outside the scenarios for which they have been trained is unclear (Lourenço et al. 2024).

Despite improvements in the efficiency and accuracy of methods for estimating haplotypes (Delaneau et al. 2019; Meier et al. 2021; Shipilina et al. 2023), in non-model species (understood as those in which, whether or not a genome has been sequenced, it is poorly annotated and has not traditionally been a model species in the pre-genomic era), haplotype-based detection methods are still not widely used. Instead, it is more common to use interpopulation methods based on detecting molecular markers with excessively high differentiation values, known as “outliers”. But even in the case of model species, the use of haplotype-based methods to detect selective sweeps presents the problem that the same genomic pattern that could be produced by a selective sweep could also be explained under different scenarios related to factors as diverse as the quality and characteristics of the sampled data, biological characteristics related to mutation and recombination rates, as well as demographic history and the effects of purifying and background selection (Johri, Aquadro, et al. 2022; Soni et al. 2023; Soni and Jensen 2024).

Part of this problem arises from the lack of knowledge of the null distribution of the statistics used, which requires simulating the neutral biological scenario. But overall, it is clear that although a statistical tool can detect a specific genomic pattern in the data, it is unlikely that that pattern could be due solely to the effect of a selective scan. It may do so in some scenarios, but not in others. Therefore, to validate a candidate SNP or region as a result of a selective process, it is first necessary to prove that the statistic does not generate false positives in realistic scenarios in terms of demography and other evolutionary parameters of interest. Subsequently, functional validation of these candidate loci will always be necessary (Johri, Aquadro, et al. 2022). This does not preclude that the development of statistical tools to detect genomic patterns that may be related to selective sweeps remains of great interest. It would also be interesting if that statistic had a known null distribution.

When studying a selective sweep, we can trace its effect over time (directional selection) or across space (divergent selection). Therefore, if we use two samples to compare the effect of the sweep, they can be separated by time or space. Detecting the footprint of natural selection in genomes in general, and specifically divergent selection, is important for studying speciation processes (Galindo et al. 2021) and climate adaptation (Folkertsma et al. 2024), but also for more immediate effects such as resistance to infections in commercially important marine species (Pampín et al. 2023; Vera et al. 2023).

In this work, I propose a statistic that uses the population stability index, also known as Jeffreys divergence, to compare the distribution of allelic classes of haplotypes (Labuda et al. 2007; Hussin et al. 2010) between two populations or samples. To develop the statistic I use the informational interpretation of the Price equation (Price 1972; Frank 2012a) defined for the haplotype allelic class (HAC hereafter) trait. The advantage of this statistic is that it follows a chi-square distribution when the null hypothesis (equal distribution of HACs among samples) is true. This not only increases computational efficiency by several orders of magnitude but also allows for the testing of biological models expected to deviate from this hypothesis, including the presence of local selection and its corresponding selective sweep. Below, I will present the development of the statistic and then demonstrate its behavior with both simulated and real genomic data from various samples of the SARS-CoV-2 virus.

## The Price equation and the population stability index for comparing population genomes

### Price equation

The Price equation in its most general formulation describes the change between two populations at any scale, spatial or temporal (Frank 2012a; Frank 2017). The equation partitions the change into a part due to natural selection and another part due to other effects. We compare two populations or frequency distributions which can be separated by space and/or time. Natural selection causes populations to accumulate information, which is measured in relation to the logarithm of biological fitness *m*= log(ω), where ω is the relative fitness (Frank 2012b; Frank 2012a).

Therefore, let *z* be a character that takes different values *z*_*i*_ with associated frequency *p*_*i*_ in population *P* and with frequency *q*_*i*_ in population *Q*. If we consider the logarithm of fitness as the character, *z* = *m*, we have that the mean change in *m* due to the effect of natural selection in one or the other population is (Frank 2012a)

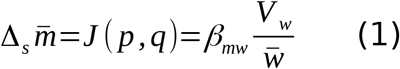

where *J* is the Jeffreys divergence or population stability index, *p* and *q* the frequency of the different values of *m* in the populations *P* and *Q* respectively, and β_*mw*_ is the regression of *m* on the absolute fitness *w*.

However, it is possible to use scales other than the fitness logarithm to measure information, with the key element being the regression of values in the new scale on fitness (Frank 2013). Therefore, to detect the effect of natural selection from genomic data, it will be necessary to measure those genomic patterns with high regression values on biological fitness. In this work, I propose using HACs as a suitable pattern to capture the increase in information generated by natural selection, whether in temporal comparisons (directional selection) or spatial comparisons (divergent selection).

### Haplotype allelic class (HAC)

HACs were initially introduced in (Labuda et al. 2007) and later used to detect genomic patterns caused by selective sweeps (Hussin et al. 2010) and divergent selection (Carvajal-Rodríguez 2017).

Consider a sample of sequences and compute the reference haplotype *R* as the one formed by the major allele of each site. Now, consider for the same or another sample of sequences, the haplotypes of length *L*+1 centered in a given candidate SNP *c* and define the mutational distance between any haplotype and the reference *R* as the Hamming distance between the haplotype and the reference i.e. the number *h* of sites in the haplotype carrying an allele different to the one in *R*. Each group of haplotypes having the same *h* will constitute a HAC (Labuda et al. 2007; Hussin et al. 2010). The HAC distribution is estimated from the distribution of the *h* values in a sample.

Thus, in a given haplotype with the candidate SNP position *c* in the middle, for each position other than *c* we count the outcome *X*_*k*_ = *I*(*s*_k_ ≠ *r*_k_) were *s*_k_ is the allele in the position *k* of the haplotype, *r*_k_ is the allele in the reference and *I*(A) is the indicator variable taking 1 if A is true and 0 otherwise. Therefore, the *h* value of an haplotype of length *L*+1 is

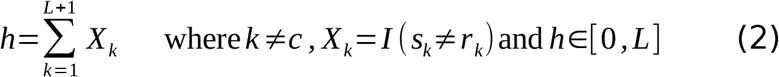

The idea behind using *h*-values to detect selective sweeps is that if one allele increases in frequency due to the effect of selection, the higher frequency alleles from adjacent sites will be swept along with the selected allele so that these haplotypes will have many common alleles with the reference configuration, i.e., an *h*-value close to zero.

### Information for HACs: the population stability index

Let *h*_*i*_ be the HAC value that satisfies *h* = *i* with *i*∈[0, *L*] then for a sample of *n*_1_ sequences in *P*, the frequency of *h*_i_ is

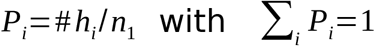

where #*h*_*i*_ is the number of occurrences of *h*_*i*_.

Similarly, for a sample of *n*_*2*_ sequences in *Q*, the frequency of *h*_i_ is

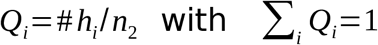

In previous works, studying the distribution of alleles around a candidate site in both samples *P* and *Q*, has been performed comparing in several ways the HAC variances of the partitions that have the reference allele or not in the different samples (Carvajal-Rodríguez 2017; Gabián et al. 2022). There are some problems with this type of approach as the unknown distribution of the defined statistics or a loss of power when using homogeneity variance tests. Here, I rely on the abstract model of the Price equation as proposed by Frank (Frank 2012a; Frank 2013; Frank 2017; Frank 2020) to calculate, using Jeffreys divergence, the change caused by selection in the distribution of HAC values between two populations.

#### Number of classes and smoothing

For a total of *L*+1 different classes the Jeffreys divergence is (Kullback 1997)

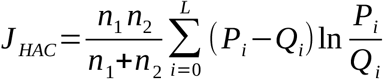

However, computing *J*_*Hac*_ in this way could suffer from the curse of dimensionality (Hastie et al. 2009) if eventually *L* > *n*_1_+*n*_2_ which will cause the presence of the different classes to be scarce. To alleviate this problem we will group the values in *K* (*K* ≤ *L* + 1) HAC classes. The number of classes *K* is an important parameter because too many classes have the dimensionality issue but too few classes will have low power for the distribution comparison. A conservative heuristic guess is *K* = (*L* +1)/2 when *L* >= 15 or *K* = *L* otherwise, since we have empirically verified that less than 15 classes implies a low detection power possibly because a smaller number of classes implies very few SNPs, which may be due to very short sequences, and/or very low sample size, and/or very homogeneous samples.

Given *K*, we will group uniformly the *h* values into *K* groups so that the first group indicates classes with equal or less than (100/*K*)% of minor alleles, the next corresponds to classes with more than (100/*K*)% but equal or less than 2×(100*/K*)%, until the last group with more than (*K*-1)×(100*/K*)% but equal or less than 100%. If necessary (*L*+1 not divisible by *K*), the class with 100% of minor alleles is included in this last group. For simplicity we consider *K* is a divisor of *L* + 1.

Thus, for population *P*, the frequency *P*’_i_ of each group of classes is

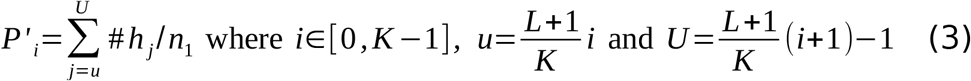

However, note that the Jeffreys divergence is defined only if *P* and *Q* have no zeros. To avoid zeros we use additive smoothing (Manning et al. 2008) with a pseudocount α=0.5 for each possible outcome so that *P’*_*i*_ in (3) become

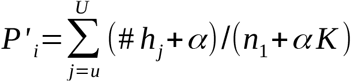

So, for *K* groups of HAC classes, the Jeffreys divergence for comparing the HAC distribution between populations *P* and *Q* finally is (c.f. eq. 5.10 in Kullback 1997 p. 130)

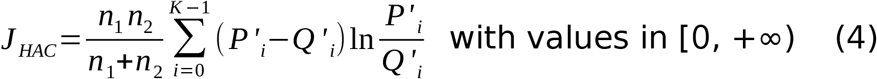

Figure 1 shows an example of a *J*_*HAC*_ calculation for two samples with 8 haplotypes each (*n*_1_ = *n*_2_ = 8). Haplotypes have a length of 9, so discounting the candidate site we have 8 sites and 9 possible HAC classes (from 0 to 8). Note that if a class does not appear in either of the two samples, the contribution value to *J*_*HAC*_ for that class is 0. When the class is present only in one sample, to correct the problem of zeros, pseudocounting is applied with α = 0.5, so that the frequency value of the class that does not appear will be 0.5/(8+9/2) = 0.5 / 12.5 = 1 / 25 in that sample. If the count in a class is 2, the corrected frequency value will be 2.5 / 12.5 = 5 / 25 (Figure 1).

**Figure 1.**
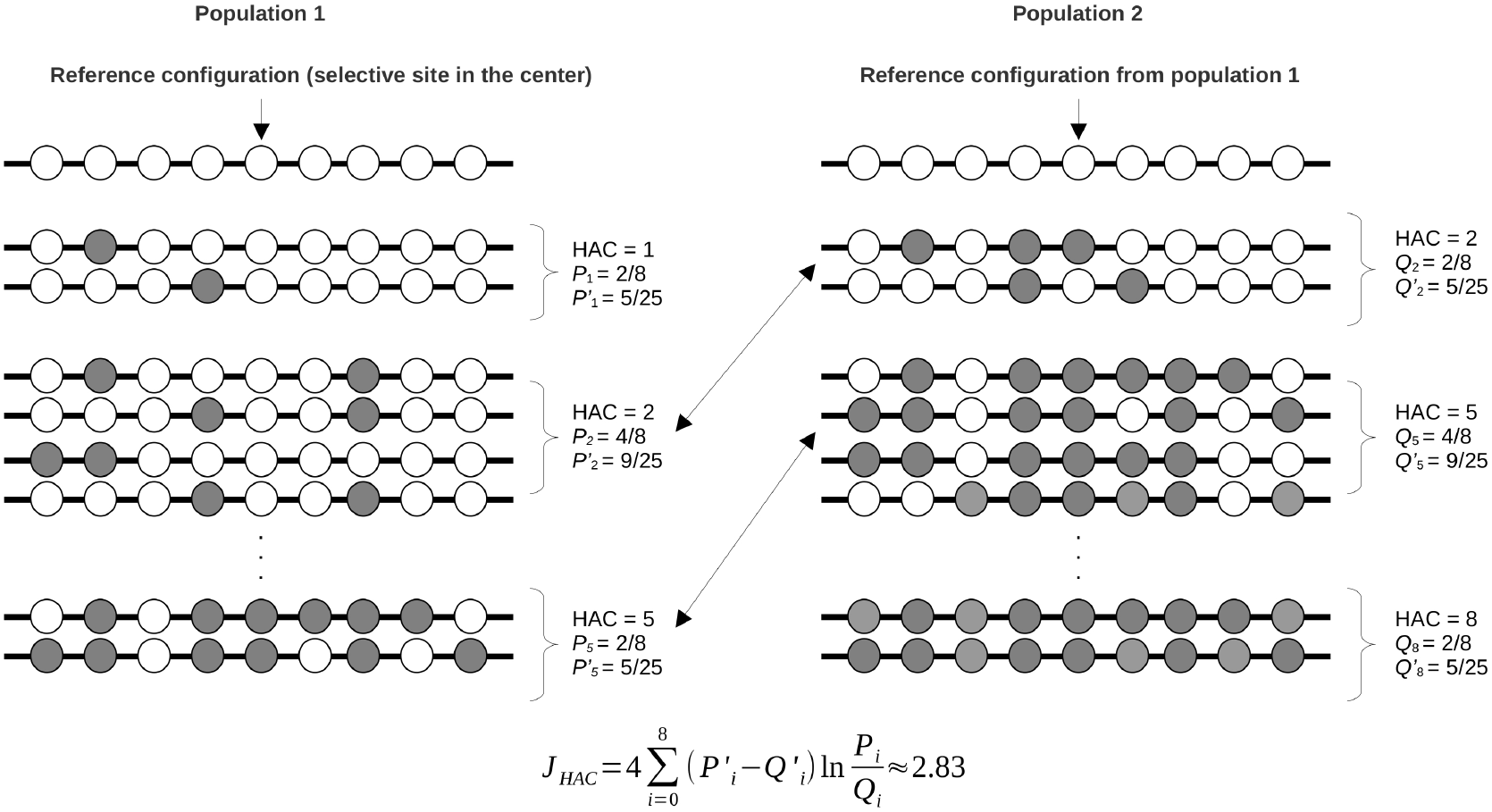
Reference configuration and the distribution of HAC values in two samples and calculation of the *J*_*HAC*_ statistic. The haplotype length is 9 and the number of haplotypes is 8 in each sample. White circles represent major alleles and grey circles represent minor alleles.

Note that *J*_*HAC*_ is also known as the population stability index and is asymptotically distributed as Chi-square with *K*-1 degrees of freedom.

The advantage of using (4) in the context of studying the genomic footprint of selection is that, contrary to other statistics, it can be approached by a chi-square distribution providing a faster approach as we can avoid performing computationally expensive simulations or resampling.

### Phenotypic scale, linkage disequilibrium and window size

#### Phenotypic scale

The gain in information caused by the effect of natural selection as expressed in (1) depends on the log-fitness *m* and if we measure the frequency of the *h*_i_ classes instead of fitness classes, the relationship between the average change in the *h* distribution and the gain in information will depend on the regression of *h*-values on fitness as follows (Frank 2013)

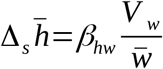

thus, if we use the HAC values to compute *J* we obtain *J*_*HAC*_

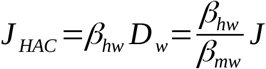

The quantity *β*_*hw*_/ *β*_*mw*_ is the change in phenotype (HAC values) relative to the change in information (Frank 2013). Therefore, if there is perfect fit between ln(*P*/*Q*) and *m* then *J*_HAC_ = *J*.

The regression of *h* on *w* will be high when it is fitness that is distributing the classes of *h*, which requires that there are indeed one or more sites under selection within the haplotype window. However, this is a necessary but not sufficient condition. Price’s equation for total change indicates that the average variation in phenotype *h* has two components: one due to selection and the other due to other causes, including changes in the components of the phenotype that are transmitted (Δ*h*)

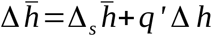

In our context, the change in *h* not caused by selection may be due to, besides mutation, the effect of recombination on haplotypes, which in turn will depend on the window size. Therefore, we are interested in using window sizes that correspond to haplotype blocks in order to minimize Δ*h*.

#### Window size

The program computes haplotype blocks and set the candidate position *c* in the middle of each block. An haplotype block is computed as a sequence of reference SNPs with length *W* that satisfies *r*^2^(*c* - *W*/2, *c* - *W*/2+1),…, *r*^2^(*c* - 1, *c*), *r*^2^(*c, c* + 1), *r*^2^(*c + x -* 1, *c* + *x*),…, *r*^2^(*c* + *W* / 2 - 1, *c* + *W* / 2),… where *r* is the correlation coefficient calculated from the sample of size *n*, so that *Pr*(*nr*^2^) ≤ α, and *nr*^2^ has a Chi square distribution. Furthermore, for a given SNP *c*+1 to be included in the block, it is also required that *D*’(*c, c*+1) ≥ 0.4, where *D*’ is the normalized linkage disequilibrium (Lewontin 1964). The block is extended until any of both conditions is rejected i.e. *Pr*(*nr*^2^_*c*+*x*-1,*c*+*x*_) > α or *D*’(*c*+*x*-1, *c*+*x*) < 0.4.

Optionally, the program can use an outlier as the putative center of a block and build the block around it. In this case, the condition for defining a block is more liberal, allowing blocks that have a mean normalized linkage disequilibrium value greater than zero. The reason is that the outliers may have been part of older blocks, so we use the minimum condition that the average linkage of reference alleles is greater than zero assuming that, if they are not the product of selective sweep, the distribution of HACs will not be affected, the latter will be checked in the next section by simulation.

### Simulations

To check the performance of the method, its power and its control of false positives, we will perform two types of simulations, namely, with diploid and haploid genomes.

#### Diploid genomes

For diploid genomes, the same simulated data as in (Carvajal-Rodríguez 2017; Gabián et al. 2022) were used. Two populations of 1000 facultative hermaphrodites were simulated under divergent selection and different conditions about mutation, recombination, migration and selection. Each individual consisted of a diploid chromosome of length 1Mb. In Tables 1-4 we can appreciate the different cases with the corresponding parameter values. The population migration rate was *Nm* = 10 in all cases. The number of generations was 10^4^ or 5×10^3^, the population mutation rate θ = 4*Nµ* was {12, 60} where *µ* is the mutation rate per haploid genome, the population recombination rate ρ = 4*Nr* was {0, 4, 12, 60} where *r* is the recombination rate per haploid genome, or noted as infinite when the segregation was independent. The selection coefficient *s* was ± 0.15 depending on if the mutant allele is deleterious (+0.15) or beneficial (-0.15).

**Table 1.** Percent power for detecting divergent selection by *J*_*Hac*_ in simulated data with the selective site in the middle. The power was computed as 100×the number of replicates where selection was detected/1000. In parentheses, the corresponding value when the blocks were built around outliers instead of finding the blocks automatically, if the value is equal the = symbol appears. Genome size is 1Mb. Population size *N*= 1000. *T*: number of generations. Population mutation rate θ = 4*Nµ*. Population recombination rate ρ = 4*Nr. s*: selection coefficient. *Dist*: average distance in Kb from the detected position to the actual effect, given only when ρ>0. *W*: average size, in number of SNPs, of the haplotypes analyzed. Significance level α = 0.05. Each case was replicated 1,000 times.

| Case | $T$ | $\theta$ | $\rho$ | $s$ | % power | Dist Kb | $W$ |
| --- | --- | --- | --- | --- | --- | --- | --- |
| C1 | $10^4$ | 12 | 0 | $\pm 0.15$ | 100 (98) | - | 14 (13) |
| C2 | $10^4$ | 12 | 4 | $\pm 0.15$ | 100 (98) | 42 (38) | 14 (13) |
| C3 | $10^4$ | 12 | 12 | $\pm 0.15$ | 100 (96) | 4 (10) | 13 (12) |
| C7 | $5 \times 10^3$ | 60 | 0 | $\pm 0.15$ | 100 (94) | - | 13 (12) |
| C8 | $5 \times 10^3$ | 60 | 4 | $\pm 0.15$ | 100 (85) | 37 (14) | 13 (12) |
| C9 | $5 \times 10^3$ | 60 | 60 | $\pm 0.15$ | 98 (80) | 14 (2) | 12 (11) |
| C13 | $10^4$ | 60 | 0 | $\pm 0.15$ | 100 (100) | - | 14 (=) |
| C14 | $10^4$ | 60 | 4 | $\pm 0.15$ | 99 (100) | 126 (15) | 14 (13) |
| C15 | $10^4$ | 60 | 60 | $\pm 0.15$ | 95 (91) | 19 (2) | 13 (12) |
| C15Indep | $10^4$ | 60 | ∞ | $\pm 0.15$ | 0 (2*) | - (-) | - (11) |

\* Note that this 2% results from using the outlier-centered haplotype method. When directly inspecting outliers with the EOS method, the power was 78%.

#### Input settings for diploid simulation data

A minor allele frequency (MAF) value of 0.01 was used. As we have already seen, the program allows defining the window or haplotypic block size automatically, using the correlation between pairs of sites to define the block size and placing the central SNP as a candidate or, alternatively, it uses the detected outliers as candidate SNPs and then calculates the window size. Both methods were used. All other parameters were as defined by default (maximum window size 1000, minimum window size 11, significance level 0.05, etc, see the program manual). An example of the command line to launch case C1 (Table 1) and analyze the 1000 files located in subfolder C1 and using the automatic calculation of blocks (-useblocks 1) is:

> *./iHDSel0.5.2 -path /home/data/C1/ -runs 1000 -input Om_SNPFile_Run -format ms -sample 50 -minwin 11 -output JHAC_C1_ -maf 0.01 -useblocks 1 -doEOS 1 &*

The -*doEOS* tag indicates whether we want (1, default) or not (0) to run in addition the EOS outlier test (Carvajal-Rodríguez 2017). If the calculation without blocks is used (-useblocks 0) the doEOS tag must necessarily be set to 1.

### Diploid Simulation results

In the following tables the results of power (Tables 1-3) and false positive rate (Table 4) after analyzing 1000 replicates of each scenario are presented. In summary, for haplotypes with linkage and the selective site in the center of the chromosome, when using the automatic blocks system, the power is equal to or greater than 95%, regardless of mutation and recombination rates. As expected, if the sites are not linked, the method does not work because there is no selective sweep (Table 1).

When the position of the selective site moves away from the center of the chromosome (Table 2), the power remains high. Localization improves as recombination increases and as the marker is located closer to the center. In the case of multiple selective sites (Table 3), the power to detect at least three is above 75% when using automatic blocks but only detects one (97% power) in the case of blocks centered on outliers. In general, for blocks centered on outliers, the power is slightly lower, but in some cases, the localization was considerably more accurate.

**Table 2.**
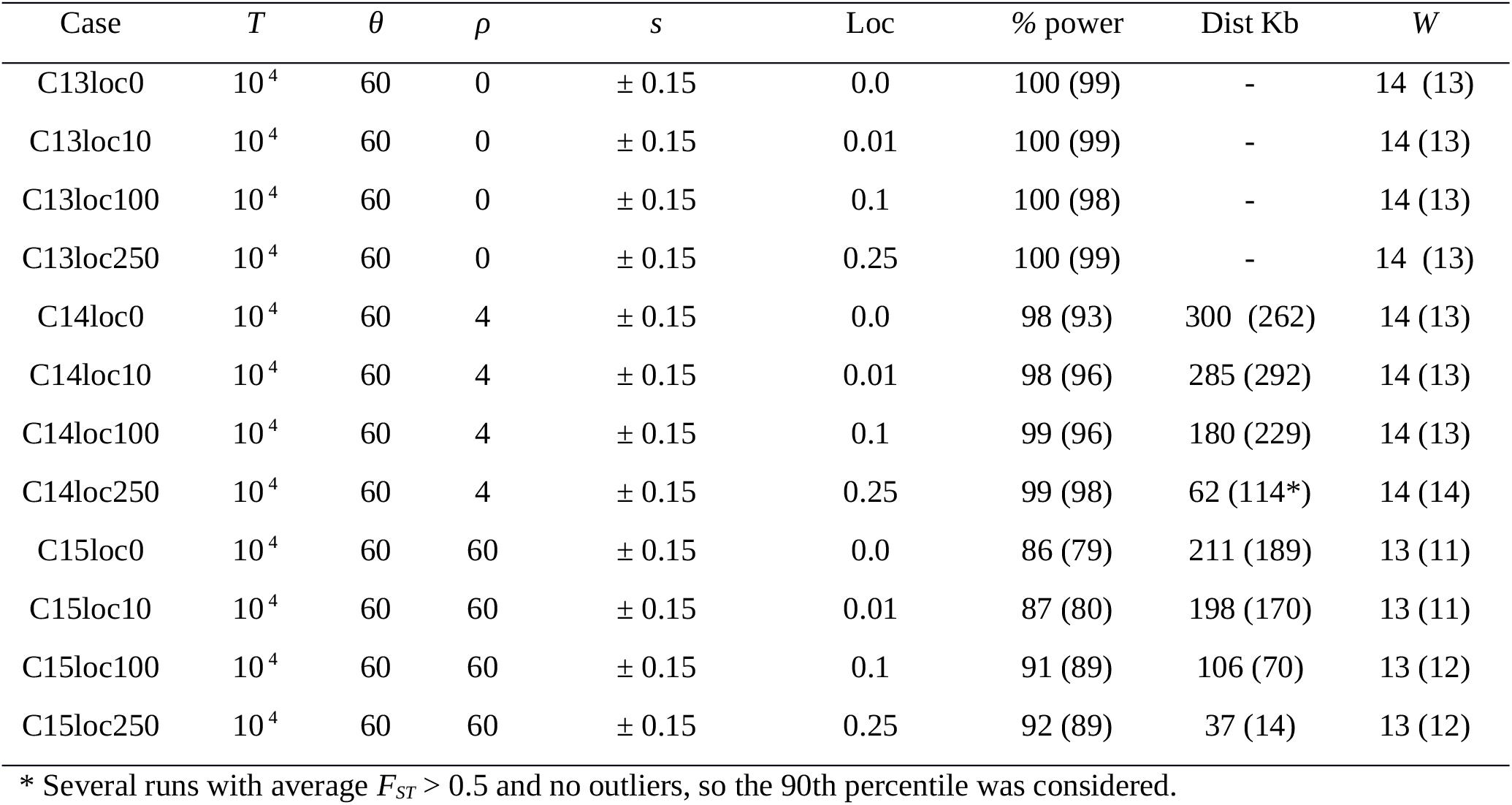
Percent power for detecting divergent selection by *J*_*Hac*_ in simulated data with the selective site in different locations. The power was computed as 100×the number of replicates where selection was detected/1000. In parentheses, the corresponding value when the blocks were built around outliers instead of finding the blocks automatically, if the value is equal the = symbol appears. Genome size is 1Mb. Population size *N*= 1000. *T*: number of generations. Population mutation rate θ = 4*Nµ*. Population recombination rate ρ = 4*Nr. s*: selection coefficient. *Loc*: true relative position of the selective site. *Dist*: average distance in Kb from the detected position to the actual effect, given only when ρ>0. *W*: average size, in number of SNPs, of the haplotypes analyzed. Significance level α = 0.05. Each case was replicated 1,000 times.

**Table 3.** Percent power for detecting divergent selection by *J*_*Hac*_ in simulated data for a polygenic model with 5 selective sites uniformly distributed in the chromosome. The power was computed as the number of replicates where selection was detected. In parentheses the corresponding % power when the blocks were built around outliers instead of finding the blocks automatically, if the value is equal the = symbol appears. Genome size is 1Mb. Population size *N*= 1000. Number of generations *T*=10^4^. Population mutation rate θ = 4*Nµ=*60. Population recombination rate ρ = 4*Nr=*60. Selection coefficient per site s=± 0.032. *W*: average size, in number of SNPs, of the haplotypes analyzed. Each case was replicated 100 times.

| Case | Candidate | % power | W |
| --- | --- | --- | --- |
| C15poly | 1 | 99 (97) | 15 (18) |
| C15poly | 2 | 89 (0) | 16 |
| C15poly | 3 | 75 (0) | 16 |
| C15poly | 4 | 59 (0) | 16 |
| C15poly | 5 | 44 (0) | 16 |

Finally, in the neutral simulations where there was no selective site (Table 4), the false positive rate conservatively remains below the expected 5%, both using automatic blocks and those centered on outliers, with one exception corresponding to the effect of bottlenecks. When a bottleneck occurs, it can generate linkage disequilibrium that could resemble the effect of a selective sweep, thus increasing the possibility of false positives (Barton 1998; Thornton and Jensen 2007; Harris et al. 2018). In our case, we observed that *J*_*Hac*_ becomes liberal with 13% when the blocks are centered around the outliers, which means an 8% excess over the expectation. The explanation for this happening with blocks centered on outliers but not with automatic ones is that, as previously indicated, the construction of blocks centered on outliers is somewhat more liberal, validating as blocks those regions that have an average disequilibrium greater than 0. A conservative option available for the above exception is to set the window size to a higher value, say 25 or 50, which solves the problem and sets the false positive rate to just 2%. While for the corresponding selective case when we run the program with these window sizes the power is 90%. The underlying logic is that if the positive is due to an increase in the frequency of a pattern randomly generated by the bottleneck, then some increase in the window size, say doubling it, will undo the effect. However, if the positive is real, it is relatively easy for it to remain unless the sweep has already been detected at the limit of its size.

**Table 4.**
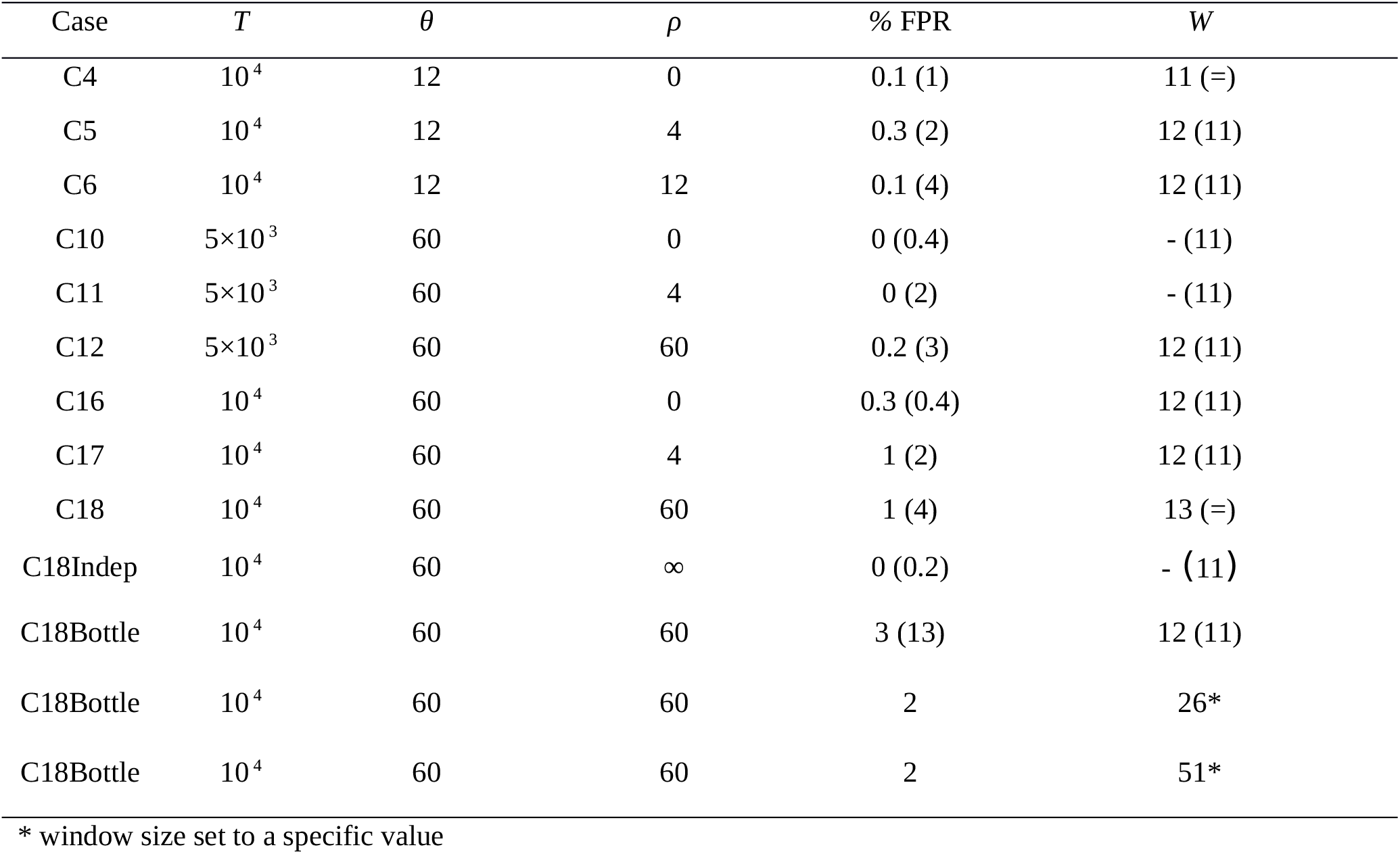
Percent false positive rate for detecting divergent selection in simulated neutral data. In parentheses the corresponding value when the blocks were built around outliers instead of finding the blocks automatically, if the value is equal the = symbol appears. Genome size is 1Mb. Population size *N*= 1000. *T*: number of generations. Population mutation rate θ = 4*Nµ*. Population recombination rate ρ = 4*Nr*. %FPR = 100×number of replicates with significant *J*_*Hac*_ test/1000. *W*: average size, in number of SNPs, of the haplotypes analyzed. Each case was replicated 1,000 times.

#### Haploid genomes

For haploid genomes, we consider a scenario related to the SARS-CoV-2 virus, which belongs to SARS (Severe Acute Respiratory Syndrome)-related viruses and is a positive-sense single-stranded RNA virus with a 30 kb genome. The model is based on average parameter values associated with the intrapatient evolutionary dynamics of the virus (Terbot, Johri, et al. 2023; Terbot, Cooper, et al. 2023).

The virus’s generation interval for one year corresponds to 861.64 viral cycles/year (J.W. Terbot et al. 2023), and the lower estimate of the sampling time per individual after infection onset is approximately 7 days, equivalent to 17 generations.

The simulated scenario corresponds to a genome of 30,000 nucleotides, an effective population size of 10,000, a mutation rate per site of 2×10^?6^, resulting in a mutation rate µ = 0.06 per genome, and a recombination rate with the same value or absent. The population starts in a mutation-drift equilibrium, where the effective number of alleles, *n*_e_ has been calculated, defining a mutant proportion of 1/ *n*_*e*_ (Crow and Kimura 1970).

Two selective sites are defined in the Spike region at nucleotide positions 23,403 and 23,604, with a favorable selection coefficient of 0.2 for the mutation. The three nucleotides corresponding to the triplets carrying these mutations (23,402-23,404 and 23,603-23,605) are set at an initial frequency of 5% at the beginning of the simulation, after reaching mutation-drift equilibrium.

The populations to be compared consist of two samples: one taken after 7 days of infection (17 generations) and another taken at the end of the simulation after an additional 7-8 days (generation 35). Each sample size is 1000. We conducted 1000 runs for each simulated case.

Two different scenarios are considered. The first scenario, without a bottleneck, includes two cases: one with selection on the two indicated sites and a neutral one. The second scenario simulates a bottleneck that occurs after 7 days and corresponds to a new infection, starting with only 1, 2 or 5 viruses. The bottleneck is modeled as discrete logistic growth (Roughgarden 1996) with a growth rate of 2 and an initial value corresponding to the founder effect (1, 2 or 5). For each of these situations, we simulate a neutral and a selective case. Simulations were carried out with a modified version of the GenomePop2 program (Carvajal-Rodriguez 2008).

Tables 5x and 6x present a summary of the parameter values used, along with the results of the detection power and false positive rate of the *J*_*Hac*_ statistic.

**Table 5.** Percent power for detecting directional selection by *J*_*Hac*_ in simulated data for haploid genomes. The number of generations was 35. Population size *N*= 30,000. Population mutation rate θ = 2*Nµ*. Population recombination rate ρ = 2*Nr. s*: selection coefficient. Bottleneck: initial bottleneck at generation 17, a dash implies that there is no bottleneck. Each case was replicated 1,000 times.

| $\theta$ | $\rho$ | $s$ | Bottleneck | % power |
| --- | --- | --- | --- | --- |
| 1200 | 0 | -0.2 | - | 66 |
| 1200 | 1200 | -0.2 | - | 64 |
| 1200 | 0 | -0.2 | 1 | 3 |
| 1200 | 1200 | -0.2 | 1 | 4 |
| 1200 | 0 | -0.2 | 2 | 38 |
| 1200 | 1200 | -0.2 | 2 | 56 |
| 1200 | 0 | -0.2 | 5 | 53 |
| 1200 | 1200 | -0.2 | 5 | 56 |

**Table 6.**
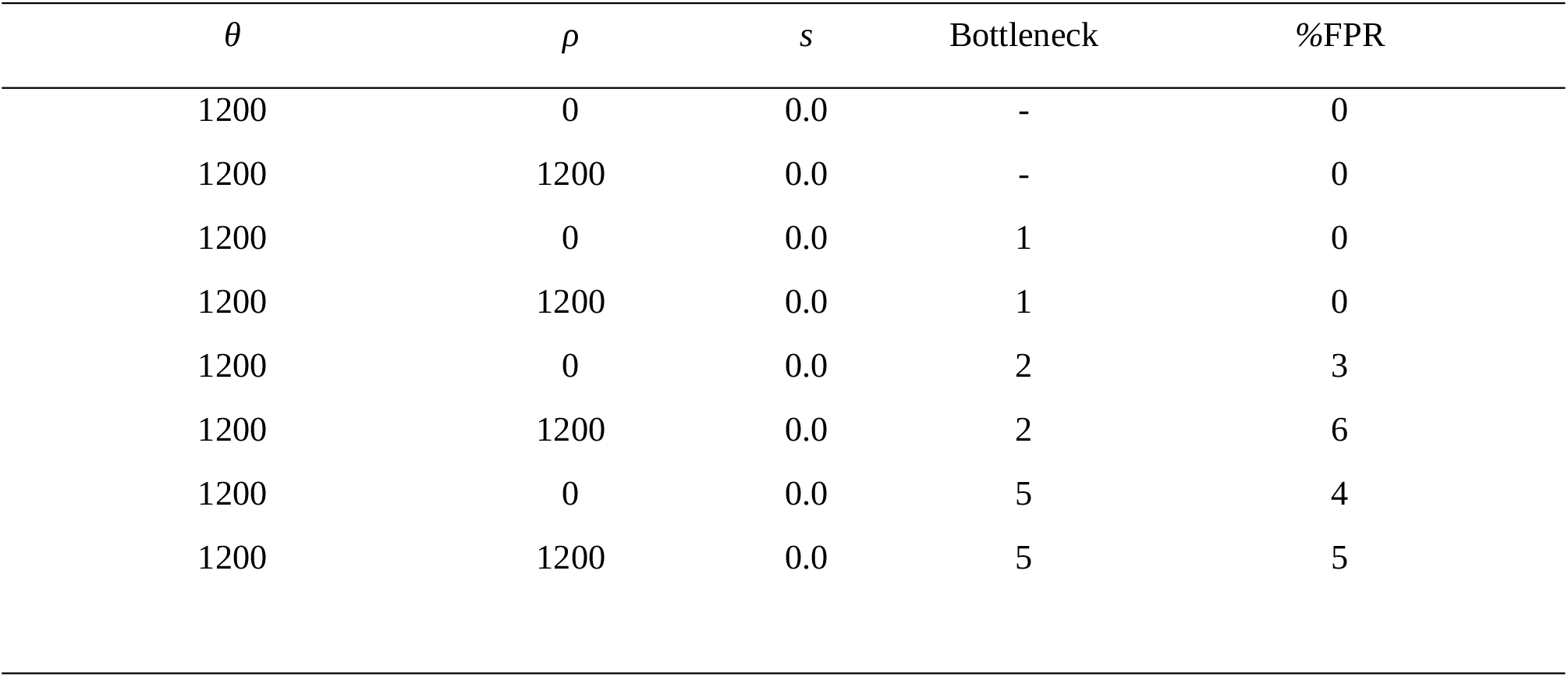
Percent false positive rate for detecting directional selection in simulated neutral data for haploid genomes. The number of generations was 35. Population size *N*= 30,000. Population mutation rate θ = 2*Nµ*. Population recombination rate ρ = 2*Nr. s*: selection coefficient. Bottleneck: initial bottleneck at generation 17, a dash implies that there is no bottleneck. %FPR = 100×number of replicates with significant *J*_*Hac*_ test/1000. Each case was replicated 1,000 times.

#### Input settings for haploid simulation data

A minor allele frequency (MAF) value of 0.01 was used. For the haploid scenario, the program was not able to detect blocks automatically possibly due to short evolutionary time and therefore we used the outliers as candidate SNPs to calculate the window size. All other parameters were as defined by default (see the program manual). An example of the command line to launch the base case and analyze the 1000 files located in subfolder Serial_M1_sites2 and using the outlier-based windows (-useblocks 0) is:

> *./iHDSel0.5.2 -path /home/data/Serial_M1_sites2/ -runs 1000 -input GP2msout_Run -format ms -sample 1000 -minwin 11 -output simSARSCv2_ -maf 0.01 -useblocks 0.*

#### Haploid Simulation results

With the simulation of haploid genomes, the power is lower due to the shorter evolutionary time but it is still around 65%. The impact of bottlenecks in this short period is to reduce the power, which is logical because it produces the loss of the selection signal. However, in the presence of recombination and with bottlenecks of two or more individuals, the power remains at 56% while the false positive rate remains between 4-6% (Tables 5-6).

### Real data analysis: SARS-CoV-2

SARS-CoV-2 virus genomes stored in the GISAID database (Khare et al. 2021) are indexed by both locality and the time period where they were sampled thus presenting a unique opportunity to apply iHDSel to both time or spatially separated samples. Therefore, as an example of application, we are going to compare SARS-CoV-2 genomes sampled in Spain (SP), England (EN) and South Africa (SA) in periods corresponding to different waves. The findings of this section are based on data associated with 30,274 SARS-CoV-2 genomes available on GISAID up to February 12, 2024, gisaid.org/EN1, gisaid.org/EN2, gisaid.org/EN3, gisaid.org/EN4, gisaid.org/SP1, gisaid.org/SP2, gisaid.org/SA.

The downloaded genomes were complete (>29,000 bp) and of high quality (<1% undefined bases and <0.05% unique amino acid mutations). These datasets were then processed using the Nextclade CLI for quality control (Aksamentov et al. 2021). Briefly, the Nextclade CLI examines the completeness, divergence, and ambiguity of bases in each genome. Only genomes considered ‘good’ by Nextclade CLI were selected.

The samples from England (EN1, EN2,EN3 and EN4) correspond to the period of March 2020, at the beginning of the first wave of the pandemic (EN1, 4820 genomes collapsed to 4227 after quality control), a second sample taken between March 28 and March 31, 2021, inclusive (EN2, 5966 genomes collapsed to 4152 after quality control), a third from June 24 to June 26, 2021, inclusive (EN3, 6886 genomes collapsed to 5844 after quality control), and from October 1, 2023, until January 31, 2024, inclusive (EN4, 3928 genomes collapsed to 3712 after quality control).

The samples from Spain (SP1 and SP2) correspond to the periods June 24, 2021, to July 12, 2021, inclusive (SP1, 6195 genomes collapsed to 4627 after quality control) and October 1, 2023, to January 31, 2024, inclusive (SP2, 1012 genomes collapsed to 221 after quality control).

Finally, the sample from South Africa corresponds to the same period as SP1, June 24, 2021 to July 12, 2021, inclusive (SA, 1467 genomes collapsed to 1327 after quality control).

These samples will allow us to compare population changes in space or time. We will compare genomes from different samples to study if there are genomic patterns that the *J*_*HAC*_ test identifies as potentially caused by selection (see below).

#### Rationale of the comparisons

##### Spatial comparisons: SP1-SA, EN3-SA, EN3-SP1

These comparisons involve samples from different countries obtained in the same time period of the pandemic. The interest in the comparison with South Africa is that on June 24, 2021 to July 12, 2021, vaccination rates were high in Spain and England but very low in South Africa. Virtually 100% of the Spanish and English population was vaccinated with at least one dose and less than 10% of the South African population (Mathieu et al. 2020).

##### Temporal comparisons: EN1-EN2, EN2-EN3, EN3-EN4

These comparisons affect the same country but in different periods of the pandemic from the beginning of the first wave to the beginning of 2024 with virtually the entire population already vaccinated several times and the majority variant being Omicron and its subvariants (Brüssow 2022; Wang et al. 2023; Wang et al. 2024).

##### Spatial comparisons: EN4-SP2

At the end of 2023, the JN.1 subvariant of Omicron, originating from the BA.2.86 lineage, began to spread. This subvariant already carried more than 30 mutations in the spike protein compared to previous subvariants.

JN.1 includes the L455S mutation and, by the end of 2023, exhibited a higher reproductive rate than previous sublineages in countries such as Spain, France, and England, with the number of detected JN.1 sequences being higher in England than in Spain (Kaku et al. 2024). During this period, DV.7.1, a sub-lineage of BA.2.75, was highly prevalent in Spain, 50% compared to 5% in the UK (O’Toole Á, Hill V, Pybus OG et al. 2021), and was considered a variant to monitor, although it was later downgraded. Therefore, the comparison between EN4 and SP2, corresponding to October 2023 - January 2024, is of interest to study the potential patterns of divergent selection in the evolutionary dynamics of Omicron subvariants between these two countries.

### Genome alignment and lineage classification

The pooled genomes for each comparison were aligned with the MAFFT FFT-NS-2 program (Katoh and Standley, 2013) with the specific version for SARS-CoV-2 accessible online (Katoh et al. 2019).

Sequences that had more than 5% ambiguous sites were removed and also, to keep the alignment length the same as the input, insertions were deleted. The remaining options were the default. After the alignment, and following the protocol recommended by NextStrain given the possibility of artifactual SNPs located at the beginning and end of the alignment (van Dorp et al. 2020), sites in the first 130 base pairs and the last 50 were removed using the program Mega X (Kumar et al. 2018). Lineages were identified with Nextclade CLI (Table 7).

**Table 7.** Percentage of SARS-CoV-2 lineages in the analyzed data.

| Data | %Alpha | %Beta | %Delta<br>(%AY.4/AY.45) | %Gamma | %Omicron<br>(%JN.1/FLIP/DV.7.1) | %Other (pre-Alpha,<br>Lambda, Mu,<br>recombinants, undefined) |
| --- | --- | --- | --- | --- | --- | --- |
| SP1 | 24 | 2 | 70 (2/0) | 2 | 0 | 2 |
| SA | 1 | 3 | 94 (0/57) | 0 | 0 | 2 |
| EN1 | 0 | 0 | 0 | 0 | 0 | 100 (pre-Alpha) |
| EN2 | 98 | 1 | 0.1 | 0.1 | 0 | 0.8 |
| EN3 | 1 | 0.02 | 98.9 (72/0) | 0 | 0 | 0.08 |
| EN4 | 0 | 0 | 0 | 0 | 96 (39/6/1) | 4 (recombinants) |
| SP2 | 0 | 0 | 0 | 0 | 97 (26/12/28) | 3 (recombinants) |

### Input settings for iHDSel

A minor allele frequency (MAF) value of 0.01 was used. The two methods already mentioned were used to define the window size (automatic or outlier-centered blocks) and the results detected by either of the two methods are reported. All other parameters were the ones by default (see program manual). An example of the command line for the comparison between EN3 and SP1 where both samples are in the file EN3_SP1.fas located in the data folder and using the outlier-centered block calculation (-useblocks 0) is:

> *-path /home/data/ -input EN3_SP1.fas -format fasta -output EN3_SP1 -useblocks 0 -tag ENGLAND &*

where -*tag* is the argument that defines the word included in the name from the England sequences and that allows to separate both samples.

Similarly, for the temporal comparison between EN2 and EN3

> *-path /home/data/ -input EN2_EN3.fas -runs 1 -format fasta -output EN2_EN3 -useblocks 0 -tag 2021-03 -reference 2*

where we have added the -*reference* tag to indicate that the EN3 sample should be used as a sample to calculate the blocks and the reference haplotype.

### The imprint of election in the SARS-CoV-2 genomes

#### Spatial comparisons: SP1-SA (summer 2021)

The SP1 sample has a majority Delta (70%) and Alpha (24%) composition while SA is mostly (94%) Delta (Table 7). The pooled SP1-SA sample consists of 247 SNPs with a frequency greater than 1%. After genome-wide analysis, iHDSel did not find any significant haplotypic blocks in the automatic search nor when focusing on outliers.

#### Spatial comparisons: EN3-SA (summer 2021)

Both samples are mostly Delta (99% EN3 and 94% SA, Table 7). The pooled EN3-SA sample consists of 107 SNPs with a frequency higher than 1%. After whole genome analysis, iHDSel found one site with the automatic block method (28282) and five sites centered on outliers (sites 7,851; 13,812; 21,846; 21,987 and 25,413 in Table 8).

**Table 8.**
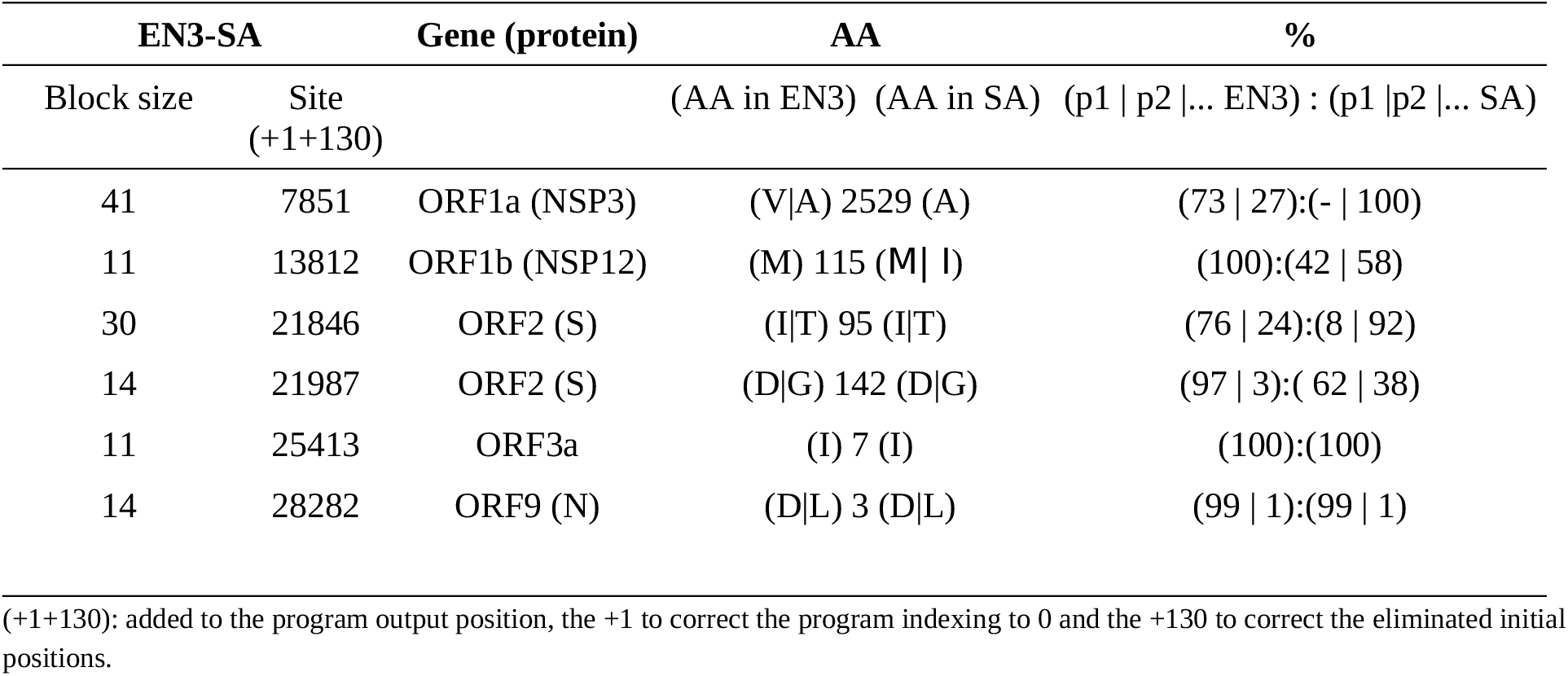
Significant *J*_*Hac*_ tests (*p-val*<0.05) for EN3-SA comparison (with 107 SNPs and sample sizes *n*_*EN3*_ = 5844, *n*_*SA*_=1327).

The first site is 7,851, which corresponds to ORF1a 2,529. In the SA sample, 100% of the sequences have the amino acid A, while in EN3, there is 27%A and 73%V, indicating the change A2529V. It is noteworthy that A2529V is one of the main SARS-CoV-2 mutations associated with virus fitness (Jankowiak et al. 2022). Moreover, in a recent study (Garcia et al. 2024) analyzing the evolution of different lineages in relation to the progress of vaccination, the A2529V mutation in ORF1a showed a significant positive correlation between the prevalence of the mutation and vaccination in Norway during the first 9 months of 2021 (including the sampling period of EN3 and SA).

The second site is 13,812, which, after identifying the slippery region (Kelly et al. 2021) and the start of ORF1b at 13,468, corresponds to amino acid 115 in ORF1b (NSP12). This site has 100%M in EN3 but 42%M and 58%I in SA. The change M115I is a characteristic mutation of the AY.45 lineage (Gangavarapu et al. 2023), which is present in SA with a frequency of 57% but is absent in EN3.

The third and fourth sites are mutations corresponding to amino acid changes in the Spike protein. Specifically, T95I represents the change observed between SA and EN3, with I at a frequency of only 8% in SA but 72% in EN3. The other mutation in Spike is G142D, with D present at 62% in SA and 97% in EN3 (Table 8). Both mutations are characteristic of the Delta variants and increase in frequency in Delta Plus (Cai and Cai 2021; Dhawan et al. 2022; Kannan et al. 2022; Mahmood et al. 2022).

The fifth site is position 25,413 of the genome, corresponding to amino acid 7 in ORF3a, with amino acid I in both samples being EN3 (ATC) and SA (ATT|50%C). Therefore, the existence of a significant signal due to different HAC distribution must be caused by accumulated variation in the surrounding sites. Similarly, the sixth and final site corresponds to amino acid 3 of the N protein, with the amino acid being D (GAT) in 99% of the cases in both samples, with practically 1% being L (CTA). Again, the existence of a significant signal due to different HAC distribution is caused by accumulated variation in the surrounding sites.

#### Spatial comparisons: EN3-SP1 (summer 2021)

We already saw that the EN3 genomes are predominantly Delta (99%), while SP1 has 70% Delta genomes and 24% Alpha (Table 7). The combined EN3-SP1 sample consists of 154 SNPs with a frequency greater than 1%. After the whole genome analysis, iHDSel found one significant site. The nucleotide site 7851 corresponds to amino acid 2,529 in ORF1a, which was also significant in the EN3-SA comparison, and we saw that A2529V is one of the main SARS-CoV-2 mutations associated with virus fitness. In this comparison, the change is from 98%A in SP1 to 73%V (27%A) in EN3.

Therefore, regarding the spatial comparisons in the summer of 2021, we see that in the SA and SP1 samples, amino acid 2529 of ORF1a was still A in virtually 100% of the sequences analyzed, while in EN3, only 27% had A and the remaining 73% were already V. This mutation is associated with an advantage for the virus and in relation to vaccination, and indeed, the *J*_*HAC*_ statistic detects it as a site with a selective pattern.

#### Temporal comparisons: EN1-EN2 (March 2020 vs March 2021)

The comparison between the English genomes is between samples separated in time (different waves). These comparisons should be considered with caution as the differentiation between samples is very large. Indeed, the mean *F*_*ST*_ in all three comparisons (EN1-EN2, EN2-EN3 and EN3-EN4) is above 0.5. However, the sites detected in the three comparisons correspond to sites with recognized impact on virus fitness.

The genomes in EN1 belong to pre-alpha variants, while the genomes in EN2 are Alpha. The combined EN1-EN2 sample consists of 77 SNPs with a frequency greater than 1%. After the whole genome analysis, iHDSel found six significant sites for the *J*_*HAC*_ test. These sites correspond to six Spike mutations, namely amino acids 501, 570, 681, 716, 982, and 1118 (Table 9). All of them correspond to the characteristic Spike mutations of Alpha (Gangavarapu et al. 2023). The only one missing is D614G, although it is included in the detected haplotypic regions. The fact that it does not come out as directly significant may be because the program did not use that position as the center of a haplotypic block, as it detected the other sites as more extreme outliers since 614G has a presence of 61%G in EN1 and 99.9% in EN2. However, when the program is run proposing the nucleotide positions corresponding to the amino acid 614 as candidates, the result is significant. Therefore, it seems that the haplotypic region including all these mutations has been detected.

**Table 9.** Significant *J*_*Hac*_ tests (*p-val*<0.05) for EN1-EN2 comparison (with 77 SNPs and sample sizes *n*_*EN1*_ = 4224, *n*_*EN2*_=4152).

| EN1-EN2 |  | Gene (protein) | AA | % |
| --- | --- | --- | --- | --- |
| Block size | Site (+1+130) |  | (AA in EN1)<br>position (AA in EN2) | (p1 \| p2 \|... EN1) : (p1 \|p2 \|... EN2) |
| 11 | 23063 | ORF2 (S) | N501Y | (100):(1 \| 99) |
| 11 | 23271 | ORF2 (S) | A570D | (100):(2 \| 98) |
| 11 | 23604 | ORF2 (S) | P681H | (100):(1 \| 99) |
| 11 | 23709 | ORF2 (S) | T716I | (100):(1\| 99) |
| 11 | 24506 | ORF2 (S) | S982A | (100):(2\| 98) |
| 11 | 24914 | ORF2 (S) | D1118H | (100):(1\| 99) |

(+1+130): added to the program output position, the +1 to correct the program indexing to 0 and the +130 to correct the eliminated initial positions.

#### Temporal comparisons: EN2-EN3 (March 2021 vs June 2021)

This is a comparison of Alpha (EN2) with Delta (EN3) genomes. The pooled EN2-EN3 sample consists of 105 SNPs with a frequency greater than 1%. After whole genome analysis, iHDSel found seven significant sites using blocks centered on outliers (Table 10).

**Table 10.**
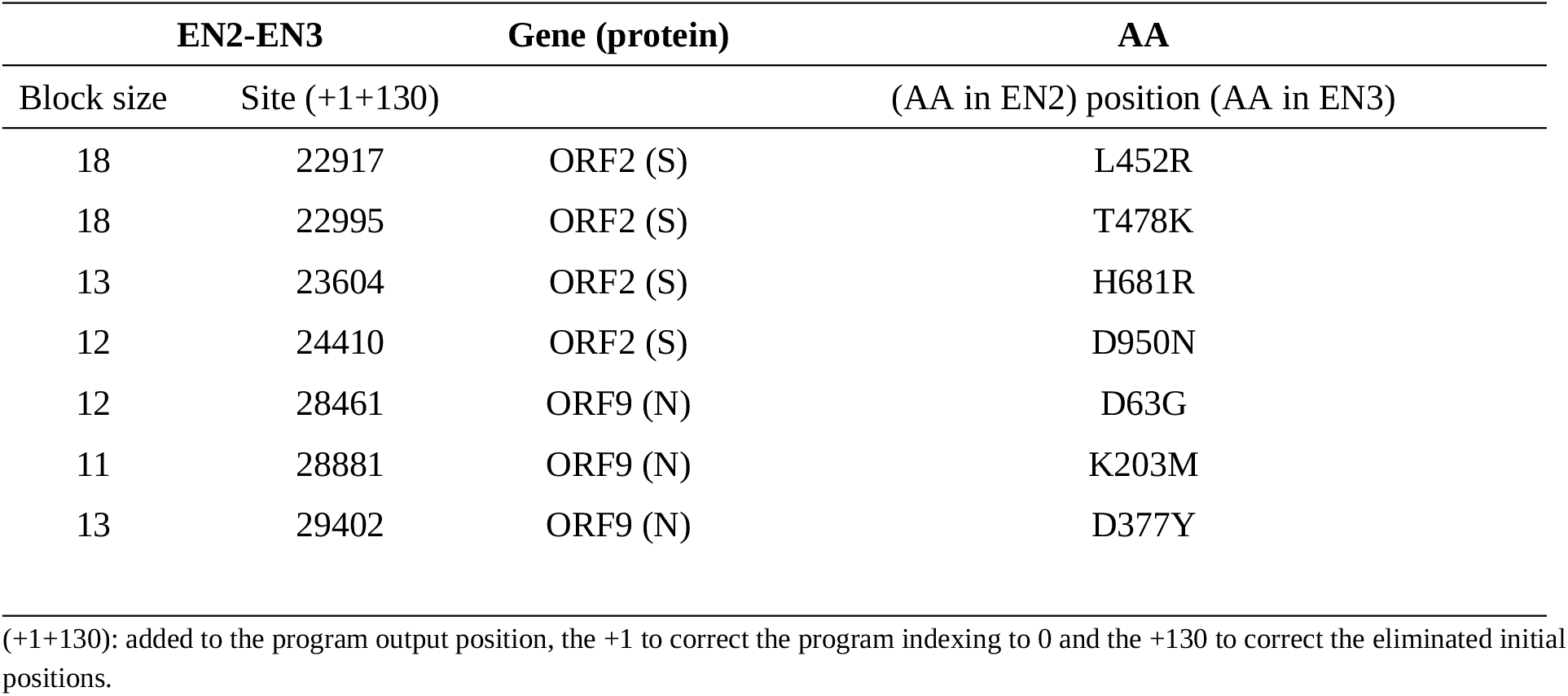
Significant *J*_*Hac*_ tests (*p-val*<0.05) for EN2-EN3 comparison (with 105 SNPs and sample sizes *n*_*EN2*_ = 4152, *n*_*EN2*_=5844).

These included substitution of relevant Spike amino acids at sites such as 452, 478, 681, and 950 (Kannan et al. 2021). For example, the L452R substitution appears to be associated with evasion of the immune response (He et al. 2022). As well as three sites in the N protein, 63, 203 and 377, which correspond to significant mutations of the delta variant, namely, D63G, R203M, and D377Y (Bhattacharya et al. 2023).

#### Temporal comparisons: EN3-EN4 (June 2021 vs January 2024)

This is a comparison of Delta genomes (EN3) with Omicron genomes (EN4). The pooled EN3-EN4 sample consists of 239 SNPs with a frequency greater than 1%. After whole genome analysis, iHDSel identified several sites with *F*_*ST*_ greater than 0.99 and 14 of them in the center of significant blocks (Table 11).

**Table 11.**
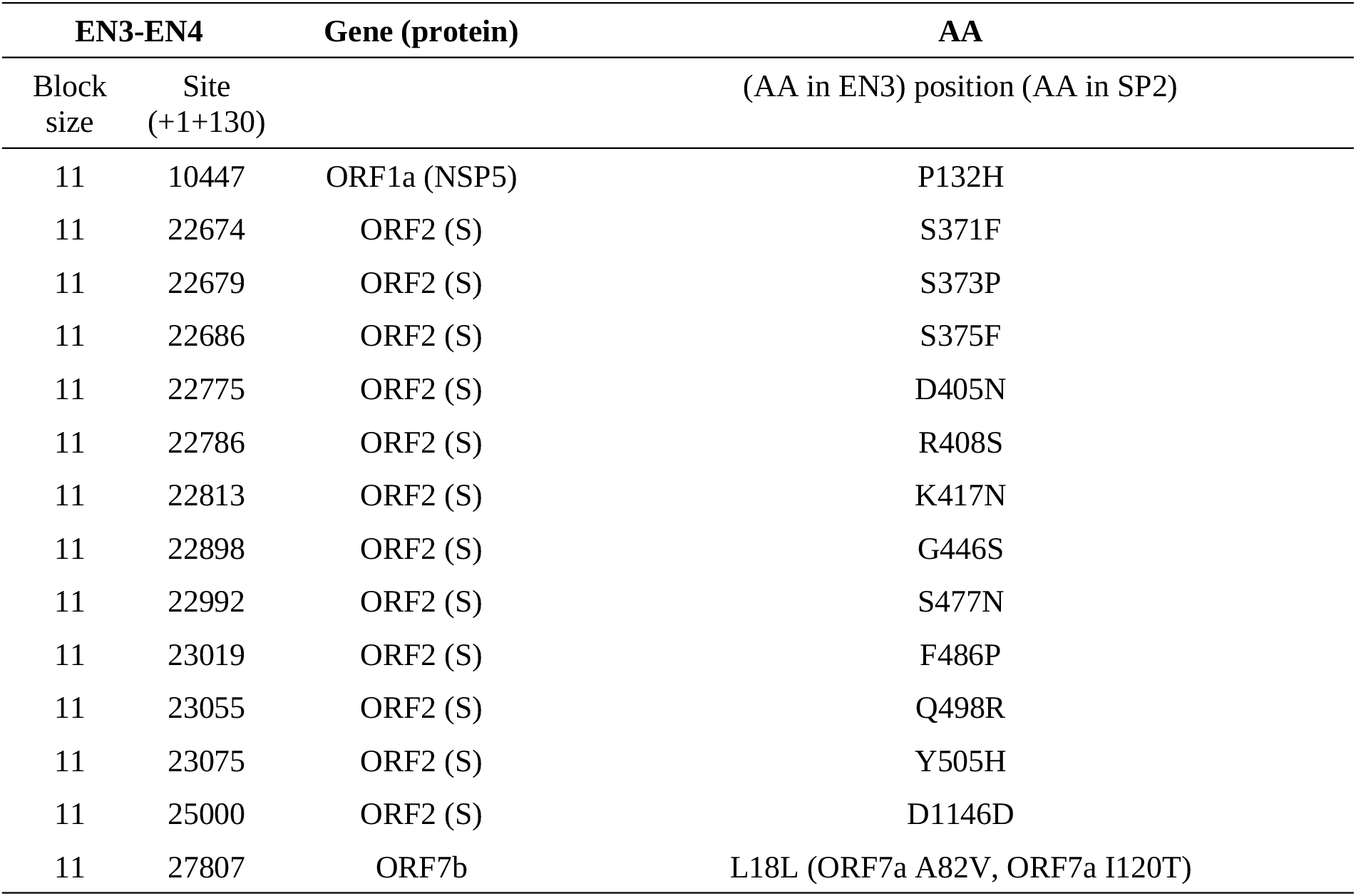
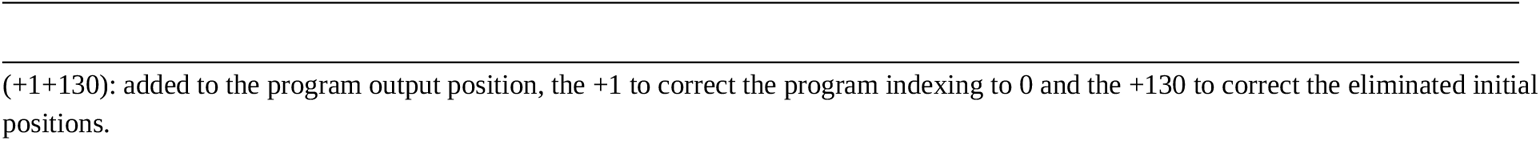
Significant *J*_*Hac*_ tests (*p-val*<0.05) for the EN3-EN4 comparison (with 239 SNPs and sample sizes *n*_*EN3*_ = 5844, *n*_*EN4*_ = 3712).

| EN3-EN4 |  | Gene (protein) | AA |
| --- | --- | --- | --- |
| Block size | Site (+1+130) |  | (AA in EN3) position (AA in SP2) |
| 11 | 10447 | ORF1a (NSP5) | P132H |
| 11 | 22674 | ORF2 (S) | S371F |
| 11 | 22679 | ORF2 (S) | S373P |
| 11 | 22686 | ORF2 (S) | S375F |
| 11 | 22775 | ORF2 (S) | D405N |
| 11 | 22786 | ORF2 (S) | R408S |
| 11 | 22813 | ORF2 (S) | K417N |
| 11 | 22898 | ORF2 (S) | G446S |
| 11 | 22992 | ORF2 (S) | S477N |
| 11 | 23019 | ORF2 (S) | F486P |
| 11 | 23055 | ORF2 (S) | Q498R |
| 11 | 23075 | ORF2 (S) | Y505H |
| 11 | 25000 | ORF2 (S) | D1146D |
| 11 | 27807 | ORF7b | L18L (ORF7a A82V, ORF7a I120T) |

(+1+130): added to the program output position, the +1 to correct the program indexing to 0 and the +130 to correct the eliminated initial positions.

The first site occurs in ORF1a (NSP5) and corresponds to the amino acid change P132H, which is a mutation in a functionally important domain and characteristic of Omicron (Hossain et al. 2022). The remaining sites presented in Table 11 correspond to core Omicron mutations in Spike (Basheer et al. 2023; Chen et al. 2023) including some like S371F, S373P, and S375F, which are related to alterations in binding and entry preference (Hu et al. 2022; Zheng et al. 2023) and also the ‘Kraken’ subvariant immune escape F486P (Parums 2023). Finally, the synonymous change L18L in ORF7b is within the same haplotypic block as the reversions A82V and I120T in ORF7a, which, when directly contrasted as candidates, were significant.

#### Spatial comparisons: EN4-SP2

The genomes of both samples are Omicrom but the subvariant composition is different (Table 7). The pooled EN4-SP2 sample consists of 218 SNPs with a frequency greater than 1%. After whole genome analysis, iHDSel identified four significant sites (Table 12).

**Table 12.**
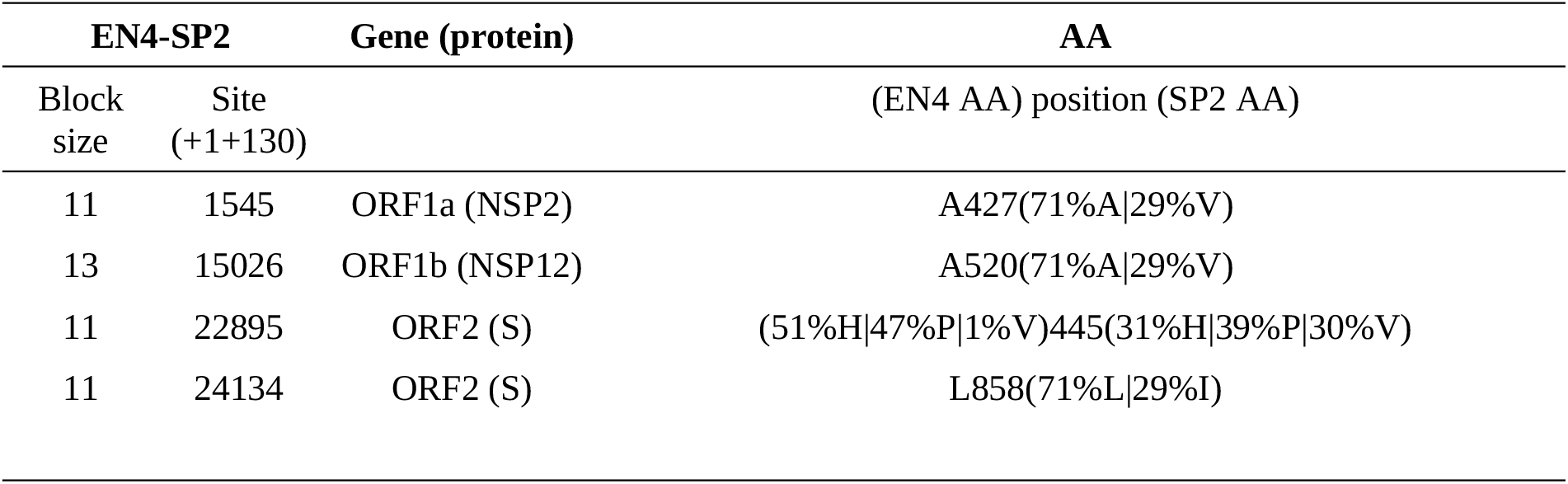
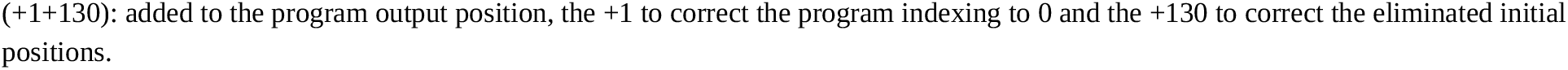
Significant *J*_*Hac*_ tests (*p-val*<0.05) for the EN4-SP2 comparison (with 218 SNPs and sample sizes *n*_*EN4*_ = 3712, *n*_*SP2*_=221). Only amino acids with a frequency equal or greater than 1% are indicated.

| EN4-SP2 |  | Gene (protein) | AA<br>(EN4 AA) position (SP2 AA) |
| --- | --- | --- | --- |
| Block size | Site (+1+130) |  |  |
| 11 | 1545 | ORF1a (NSP2) | A427(71%A\|29%V) |
| 13 | 15026 | ORF1b (NSP12) | A520(71%A\|29%V) |
| 11 | 22895 | ORF2 (S) | (51%H\|47%P\|1%V)445(31%H\|39%P\|30%V) |
| 11 | 24134 | ORF2 (S) | L858(71%L\|29%I) |

(+1+130): added to the program output position, the +1 to correct the program indexing to 0 and the +130 to correct the eliminated initial positions.

The change A427V in ORF1a is characteristic mutation of the DV.7.1 Omicron sublineage (Gangavarapu et al. 2023) which is virtually absent in EN4 (0.6%) but has a 28% in SP2 (Table 7) which explains the absence of 427V in EN4 and the 29%V in SP2. The same scenario applies to A520V in ORF1b. The other two significant sites belong to Spike. The mutation at 445 would be related to the V445H and V445P changes that seem to favor immune evasion of the virus (Ao et al. 2023; Chen et al. 2023) with the presence of 445V being 30% in SP2 but only 1% in EN4 (Table 12). Finally, L858I is also a characteristic mutation of DV.7.1.

## Discussion

In this work, a new statistic called *J*_*Hac*_ is proposed to detect genomic patterns compatible with selective sweeps. The statistic is constructed from the interpretation in terms of information of the Price equation (Price 1972; Frank 2012a) and consists of the population stability index applied to the distribution of haplotype allelic classes (HACs) in two samples. The iHDSel program incorporates the statistic along with the calculation of haplotype blocks in such a way that each candidate site is located in the center of a block. *J*_*Hac*_ appears to work optimally with simulated data where two diploid populations are subjected to divergent selection under different mutation and recombination conditions. However, if using the program mode that places the outlier sites in the center of the blocks, care must be taken because the false positive rate increases in bottleneck scenarios. A possible correction in these scenarios is to repeat the calculation with a slightly larger window size.

Real SARS-CoV-2 data have also been used to test *J*_*Hac*_ in both spatial and temporal comparisons. Some sites known to impact virus fitness and its ability to promote immune escape have been detected.

### The Price equation for comparing genomic patterns

The general formulation of the Price equation describes a change between two populations at any scale, spatial or temporal (Frank 2017). The Price equation has been proposed as a unifying principle in evolutionary biology, allowing the formulation and systematization of different evolutionary models and motivating the development of equations and models that reveal invariances and general principles (Luque 2017; Luque and Baravalle 2021). Here, we have used the selective component of the Price equation, specifically its interpretation in terms of information theory (Frank 2012a), which allows the expression of the covariance between fitness and the trait under study in terms of Jeffreys divergence or population stability index. We have defined as a trait the haplotype allelic class and used Jeffreys divergence to compare the distribution of the trait between two populations. The change in trait distribution would be compatible with the effect of selective sweeps, whether due to divergent or directional selection, depending on whether we are comparing populations in space or time.

### Limitations of the *J*_*Hac*_ method

The detection of selective sweeps is affected by different evolutionary and demographic scenarios. Throughout the space of the various parameters (mutation, recombination, background and deleterious selection, etc.) it is not difficult to find scenarios that generate an excess of false positives (Johri, Aquadro, et al. 2022; Soni et al. 2023). In our case, we have seen that some evolutionary scenarios, such as bottlenecks, can generate interpopulation genomic patterns that increase the false positive rate when using automatic window sizes centered on outliers. Although increasing the window size restores control over the false positive rate, it is possible that other scenarios without positive selection could also alter the HAC patterns. Furthermore, an excessively large window size will cause the recombination effect to dilute the swept signal.

Moreover, as we have already indicated, the method proposed here arises from the informational interpretation of the selective component of the Price equation. However, it is a statistical decomposition based on covariance, and we know that correlation does not imply causation. There is also no a priori guarantee that the partition between selection and transmission is additive (Okasha and Otsuka 2020). Therefore, *J*_*Hac*_ is an indirect method that detects a genomic pattern possibly related to selection but which can also be generated under other circumstances. Hence, the detected sites should be verified through direct methods such as the study of gene function, fitness, etc.

Finally, some genomic patterns of selection correlate with environmental variables, making it difficult to separate both effects (Folkertsma et al. 2024). The method proposed here could be combined with other methods that take this correlation into account.

### Concluding remarks

There are many statistics for identifying regions of selective sweeps in genomes, see for example (Horscroft et al. 2019; Stephan 2019; Horscroft et al. 2020; Abondio et al. 2022; Panigrahi et al. 2023). The use of machine learning-based methods to detect selection patterns has been increasing due to their accuracy and ability to handle large amounts of complex data. The underlying idea of all these methods is to use classification algorithms trained with known response data (simulations). That is, if we aim to detect a selection pattern, we train the algorithm with data that we know contains that pattern and with other data without the pattern. Different types of algorithms have been applied: neural networks, extremely randomized trees, and boosting algorithms (Horscroft et al. 2019; Panigrahi et al. 2023). A major advantage of these methods is their power and flexibility, partly due to the ease of incorporating new statistics with minimal changes to the structure of the method. Two recent machine learning methods have been designed to detect genomic signatures caused by natural selection, using a supervised multi-statistic machine learning approach (Arnab et al. 2023; Lauterbur et al. 2023). In this work, we have developed a new statistic, *J*_*Hac*_, which, due to its known null distribution, allows us to efficiently and quite accurately test for the existence of genomic patterns compatible with selective sweeps. Therefore, *J*_*Hac*_ could be an additional measure to consider for future AI-based selection detection methods. In addition, *J*_*Hac*_ has been incorporated into the iHDSel program (https://acraaj.webs.uvigo.es/iHDSel.html) along with an automatic haplotype block detection system, so it can be run independently or in conjunction with the heuristic EOS outlier detection method (Carvajal-Rodríguez 2017).

## Acknowledgements

This work was supported by Xunta de Galicia (Grupo de Referencia Competitiva, ED431C 2024/22), Ministerio de Ciencia e Innovación (PID2022-137935NB-I00), Centro singular de investigación de Galicia accreditation 2024-2027 (ED431G 2023/07) and ERDF A way of making Europe, and the Marine Science Programme (ThinkInAzul) supported by the Ministerio de Ciencia e Innovación and Xunta de Galicia with funding from the European Union NextGenerationEU (PRTR-C17.I1) and European Maritime and Fisheries Fund. Funding for open access charge: Universidade de Vigo/CISUG.

I gratefully acknowledge all data contributors, i.e., the Authors and their Originating laboratories responsible for obtaining the specimens, and their Submitting laboratories for generating the genetic sequence and metadata and sharing via the GISAID Initiative, on which the real data example in this article is based.

